## Supplementary Materials for "Directed cortico-limbic dialogue in the human brain"

### Supplementary Data
### Directed cortico-limbic dialogue in the human brain

This supplemental data provides additional information on the study of signal flow that was too detailed to be included in the main text and figures. It covers the re-analysis of an existing dataset to extract a cortical parcellation, supplementary methods and supplementary results, including control, correlation and sensitivity analyses.

#### Cortical parcellation: Re-analysis of the F-tract dataset

Circumvolved cortical areas are classically grouped into brain regions based on anatomical landmarks, such as gyri and sulci captured in the Destrieux's Atlas (Supplementary Table 1)[1]. Whether this conventional anatomy delineates areas of the cortex that are effectively connected is an open issue. Evidence for an effective connection from cortical area A to cortical area B results from the non-zero average response signal in B, upon repeated stimulation in A. The openly available and most complete stimulation-based brain atlas called 'F-tract' reports the incidence (%) of effective cortical connections across 583 participants among patients with epilepsy evaluated at French hospitals[2–5]. It covers all possible pairs of 77 Destrieux's cortical areas (Hippocampus and amygdala added). From this dataset, we grouped pairs of cortical areas into a topology of 10 effectively and densely connected brain regions using the Louvain algorithm, an unsupervised clustering method that identifies 'communities' in weighted and directed networks. Only slightly departing from conventional anatomical organization, cortical areas were grouped into nine (sub-)lobar regions (neocortex, thereafter) and one mesio-temporal region of limbic structures. Highlighting the known differential connectivity within the insula[6–8], we found an operculo-insular region densely connecting the frontal operculum with the anterior insula[6,9,10] and an superotemporal region connecting the posterior insula, the planum temporale and the superior temporal gyrus.[9,10]

Supplementary Table 1 shows the correspondence between conventional lobes (first column), brain regions (column 2) of effectively connected cortical areas (column 3) and Destrieux's labels (column 4). Names of cortical areas that are bolded correspond to the terminologia anatomical. Other names are the usual terminology used in clinical context. Cortical areas that were added to a cluster include the precuneus and cuneus, as they were not part of the F-tract dataset. Cortical areas that were re-assigned to a cluster include the singled out superior temporal sulcus, and the split cingulate cortex, to keep it as one region. Given the focus of this study, one brain region that were kept as separate areas/structures include the hippocampus (anterior and posterior), the amygdala and connected parahippocampal and temporo-polar cortex.

| Lobes | Effectively connected | Cortical areas | Destrieux name | Comment |
|---|---|---|---|---|
| Orbitofrontal | Orbitofrontal | Frontopolar gyri and sulci | G_and_S_transv_frontopol | |
| Orbitofrontal | | Frontomarginal gyrus and sulcus | G_and_S_frontomargin | |
| Orbitofrontal | | **Orbital gyri** | G_orbital | |
| Orbitofrontal | | **Orbital sulci** | S_orbital-H_Shaped | |
| Orbitofrontal | | **Olfactory sulcus** | S_orbital_med-olfact | |
| Orbitofrontal | | Gyrus Rectus | G_rectus | |
| Orbitofrontal | | Suborbital sulcus | S_suborbital | |
| Orbitofrontal | | **Subcallosal gyrus** | G_subcallosal | |
| Orbitofrontal | Operculo-insular | Lateral orbital sulci | S_orbital_lateral | |
| Dorsofrontal | | **Pars orbitalis** | G_front_inf-Orbital | |
| Dorsofrontal | | **Lateral fissure, antero-horizontal segment** | Lat_Fis-ant-Horizont | |
| Dorsofrontal | | **Pars triangularis** | G_front_inf-Triangul | |
| Dorsofrontal | | **Lateral fissure, antero-vertical segment** | Lat_Fis-ant-Vertical | |
| Insula | | **Circular sulcus of the insula, anterior** | S_circular_insula_ant | |
| Insula | | **Short insular gyri** | G_insular_short | |
| Insula | | **Circular sulcus of the insula, superior** | S_circular_insula_sup | |
| Dorsofrontal | | **Pars opercularis** | G_front_inf-Opercular | |
| Central | Dorsofrontal | **Precentral sulcus,** inferior | S_precentral-inf-part | |
| Dorsofrontal | | **Inferior frontal sulcus** | S_front_inf | |
| Dorsofrontal | | **Middle frontal gyrus** | G_front_middle | |
| Dorsofrontal | | Middle frontal sulcus | S_front_middle | |
| Dorsofrontal | | **Superior frontal sulcus** | S_front_sup | |
| Dorsofrontal | Central | **Superior frontal gyrus** | G_front_sup | |
| Central | | **Precentral sulcus,** superior | S_precentral-sup-part | |
| Central | | **Precentral gyrus** | G_precentral | |
| Central | | **Central Sulcus** | S_central | |
| Central | | **Postcentral gyrus** | G_postcentral | |
| Central | | **Postcentral sulcus** | S_postcentral | |
| Central | | **Paracentral gyrus and sulcus** | G_and_S_paracentral | |
| Parietal | | **Precuneus** | G_precuneus | added |
| Parietal | | **Superior parietal lobule** | G_parietal_sup | |
| Cingulate | Cingulate | Pericallosal sulcus | S_pericallosal | |
| Cingulate | | Cingulate Gyrus, Anterior | G_and_S_cingul-Ant | re-assigned |
| Cingulate | | **Cingulate gyrus, middle-anterior** | G_and_S_cingul-Mid-Ant | |
| Cingulate | | **Cingulate gyrus, middle-posterior** | G_and_S_cingul-Mid-Post | re-assigned |
| Cingulate | | Cingulate sulcus, marginal branch | S_cingul-Marginalis | re-assigned |
| Parietal | | **Subparietal sulcus** | S_subparietal | |
| Cingulate | | **Cingulate gyrus, postero-dorsal** | G_cingul-Post-dorsal | |
| Cingulate | | **Cingulate gyrus, postero-ventral** | G_cingul-Post-ventral | |
| Parietal | Parietal | **Intraparietal sulcus** | S_intrapariet_and_P_trans | |
| Parietal | | Sulcus of Jensen | S_interm_prim-Jensen | |
| Parietal | | **Supramarginal gyrus** | G_pariet_inf-Supramar | |
| Parietal | | **Angular gyrus** | G_pariet_inf-Angular | |
| Laterotemporal | Insulo-temporal | **Lateral fissure, posterior** | Lat_Fis-post | |
| Central | | Subcentral gyrus and sulcus | G_and_S_subcentral | |
| Insula | | **Long insular gyri and central sulcus of the insula** | G_Ins_lg_and_S_cent_ins | |
| Insula | | **Circular sulcus of the insula, inferior** | S_circular_insula_inf | |
| Laterotemporal | | **Transverse temporal sulcus** | S_temporal_transverse | |
| Laterotemporal | | **Planum temporale** | G_temp_sup-Plan_tempo | |
| Laterotemporal | | Gyrus of Heschl | G_temp_sup-G_T_transv | |
| Laterotemporal | | **Superior temporal gyrus** (T1) | G_temp_sup-Lateral | |
| Laterotemporal | | **Superior temporal Sulcus** | S_temporal_sup | re-assigned |
| Laterotemporal | Inferotemporal | **Middle temporal gyrus** (T2) | G_temporal_middle | |
| Laterotemporal | | **Inferior temporal sulcus** | S_temporal_inf | |
| Laterotemporal | | **Inferior temporal gyrus** (T3) | G_temporal_inf | |
| Basotemporal | | Fusiform gyrus (T4) | G_oc-temp_lat-fusifor | |
| Basotemporal | | **Lateral occipito-temporal sulcus** | S_oc-temp_lat | |
| Occipital | Occipital | Anterior occipital sulcus | S_occipital_ant | |
| Occipital | | Inferior occipital gyrus (O3) and sulcus | G_and_S_occipital_inf | |
| Occipital | | Middle occipital and lunatus sulcus | S_oc_middle_and_Lunatus | |
| Occipital | | Superior and **transverse occipital sulcus** | S_oc_sup_and_transversal | |
| Occipital | | **Occipital pole** | Pole_occipital | |
| Occipital | | Middle occipital gyrus (O2) | G_occipital_middle | |
| Occipital | | Superior occipital gyrus (O3) | G_occipital_sup | |
| Occipital | | **Parieto-occipital sulcus** | S_parieto_occipital | |
| Occipital | | **Calcarine sulcus** | S_calcarine | |
| Occipital | | **Cuneus** | G_cuneus | added |
| Basotemporal | | **Collateral sulcus,** Medial occipito-temporal | S_oc-temp_med_and_Lingual | |
| Basotemporal | | **Lingual gyrus** | G_oc-temp_med-Lingual | |
| Basotemporal | | **Collateral sulcus**, posterior transverse | S_collat_transv_post | |
| Basotemporal | Limbic | **Collateral sulcus**, anterior transverse | S_collat_transv_ant | |
| Basotemporal | | **Parahippocampal gyrus** (T5) | G_oc-temp_med-Parahip | |
| Hippocampus | | Hippocampus, posterior | PosteroHippocampus | separate |
| Hippocampus | | Hippocampus, anterior | AnteroHippocampus | separate |
| Amygdala | | Amygdala | Amygdala | separate |
| Basotemporal | | **Temporal pole** | Pole_temporal | |
| Laterotemporal | | Planum polare | G_temp_sup-Plan_polar | |

**Supplementary Table 1 | Parcellation of brain regions according to their effective connectivity.**

### Supplementary methods

#### Inselspital Participants

Participants in our study all underwent a presurgical workup for pharmacoresistant epilepsy at Inselspital, University Hospital Bern. This clinical evaluation typically lasts 5 to 12 days and aims at localizing the epileptogenic zone (EZ) based on recorded interictal and ictal (seizures) epileptiform. Supplementary Table 2 summarizes the participant, recording and epilepsy characteristics.

| Participant | | | Coverage | | | | | | | | | | | | Recordings | | | | Epilepsy | |
|---|---|---|---|---|---|---|---|---|---|---|---|---|---|---|---|---|---|---|---|---|
| | | | # Electrodes | Hippocampus | Amygdala | Basotemporal | Laterotemporal | Insular | Orbitofrontal | Dorsofrontal | Cingular | Central | Parietal | Occipital | Duration | % Asleep | Medication | Closest seizure | Localisation | Etiology |
| 1 | 45F | Working | 6 | ✓ | - | - | ✓ | ✓ | ✓ | ✓ | - | ✓ | - | - | 21h | 37% | Being reduced | 5h later | L Mesiotemporal | DNET |
| 2 | 20M | Student | 7 | ✓ | ✓ | ✓ | ✓ | ✓ | - | ✓ | - | ✓ | - | - | 23h + 46h | 23% | Being reduced | 8d later | L Mesiotemporal | Hippocampal sclerosis |
| 3 | 56F | Unemployed | 8 | ✓ | ✓ | ✓ | ✓ | ✓ | ✓ | - | - | - | - | - | 14h | 3% | Being reduced | 3h earlier | Unkown | Unkown, non-lesional |
| 4 | 33M | Working | 9 | ✓ | - | - | ✓ | ✓ | - | ✓ | - | ✓ | ✓ | - | 35h | 9% | Being reduced | 1h later | L Central | Post-ischemic |
| 5 | 42M | Working | 5 | ✓ | ✓ | - | ✓ | ✓ | - | - | - | - | - | - | 40h | 23% | Being reduced | During | Bi-mesiotemporal | Hippocampal sclerosis |
| 6 | 50M | Working | 6 | ✓ | ✓ | ✓ | ✓ | - | - | - | - | - | - | - | 49h | 34% | Being reduced | During | L Mesiotemporal | Hippocampal sclerosis |
| 7 | 58F | Working | 9 | ✓ | ✓ | - | ✓ | ✓ | - | - | - | - | ✓ | ✓ | 56h | 25% | Stopped | 5h later | L Latero-mesiotemporal | Post-ischemic |
| 8 | 60F | Working | 8 | ✓ | ✓ | - | ✓ | ✓ | ✓ | ✓ | - | - | - | - | 45h | 23% | Being reduced | 1h later | L Mesiotemporal | Hippocampal sclerosis |
| 9 | 24F | Protected workshop | 13 | ✓ | ✓ | ✓ | ✓ | ✓ | ✓ | ✓ | - | ✓ | ✓ | - | 53h | 39% | Being reduced | 3h earlier | Unknown | Unknown, non-lesional |
| 10 | 63F | Working | 11 | ✓ | ✓ | ✓ | ✓ | ✓ | ✓ | - | - | - | - | ✓ | 70h | 20% | Being reduced | 4h earlier | R Insulo-temporal | Unknown, non-lesional |
| 11 | 26M | Working | 10 | ✓ | ✓ | ✓ | ✓ | ✓ | ✓ | ✓ | ✓ | - | - | ✓ | 60h | 40% | Being reduced | 3h earlier | Bi-mesiotemporal | Unknown, non-lesional |
| 12 | 57F | Working | 10 | ✓ | ✓ | ✓ | ✓ | ✓ | ✓ | ✓ | ✓ | - | ✓ | - | 37h | 39% | Being reduced | 1h later | R Mesiotemporal | Unknown, non-lesional |
| 13 | 28M | Working | 10 | ✓ | ✓ | ✓ | ✓ | ✓ | ✓ | ✓ | - | - | ✓ | ✓ | 22h | 33% | Being reduced | 17h later | L Laterotemporal | Vascular lesion |
| 14 | 32F | Protected workshop | 11 | ✓ | ✓ | - | ✓ | ✓ | ✓ | ✓ | ✓ | - | - | - | 66h | 45% | Being reduced | During | Bi-mesiotemporal | Unknown, non-lesional |
| 15 | 24F | Working | 10 | ✓ | ✓ | ✓ | ✓ | ✓ | ✓ | ✓ | ✓ | - | - | - | 55h | 41% | Stopped | Days later | R Mesiotemporal | Hippocampal sclerosis |

**Supplementary Table 2 – Inselspital participants.** Participant, recordings and epilepsy characteristics. L: left, R: right. NA: sufficient sleep data was not available in participants 3 and 4.

##### Sleep scoring

Sleep scoring was done by two independent visual scorers (EVM and PN) based on 30-second segments of concomitant scalp EEG recordings according to the rules of the American Academy of Sleep Medicine (AASM) into one of five brain states (awake, NREM sleep stages N1, N2, N3, and REM sleep). Scoring was validated with an average preset Cohen's inter-scorer agreement > 0.70 between two independent scorers (EVM and PN).

#### Recording and stimulation

We used two CE-labeled devices from Inomed Medizintechnik GmbH, Germany, for our stimulation setup: the ISIS Neurostimulator and the ISIS Switchmatrix. The neurostimulator is engineered to deliver current-controlled electrical pulses for stimulating nerves or the brain. The Switchmatrix, connected to the neurostimulator, is a dynamic control center, which enables seamless and fast selection of the contacts for stimulation without requiring manual wiring adjustment. It directs the neurostimulator's pulses by switching pre-selected pairs of electrode contacts from recording to stimulating mode and back. The recordings of all non-stimulated iEEG contacts are routed through the switching matrix to the amplifier. This mechanism enables continuous monitoring of other electrode contacts throughout the stimulation, but blanks recording in the stimulating pair for about one second. Both devices are powered via USB cables (USB 3.0) connected to the research computer, allowing communication with the devices through a Dynamic Link Library (DLL). A BNC connector (Bayonet Neill–Concelman) connects the neurostimulator to the clinical EEG amplifier, sending triggers at stimulation onset, which is stored as a TTL (Transistor–transistor logic) signal (Supplementary Fig. 1a).

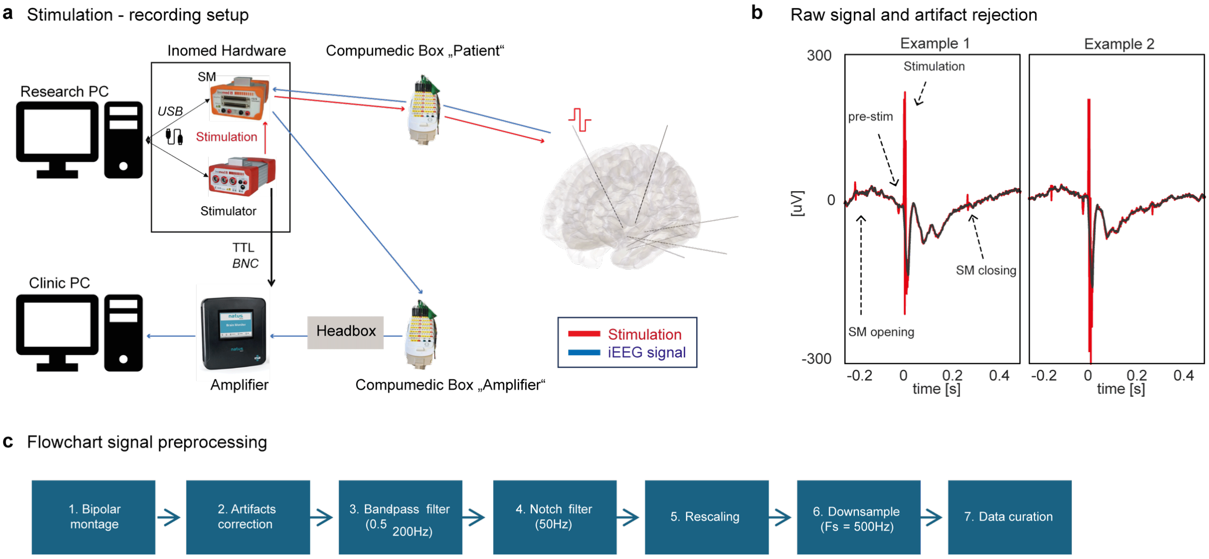


**Supplementary Figure 1 | Neurostimulation and recorded cortical responses. a**. illustration of the clinical neurostimulation-recording setup used in the study. **Red**: Electrical stimulation path. The neurostimulator sends electrical pulses to the switch-matrix (SM). The SM has previously received a command to connect two pre-selected electrodes to the neurostimulator for bipolar pulse delivery under current control. **Blue**: iEEG signal recording. Only the two pre-selected electrodes are switched off the recording chain during the time of stimulation (<1s). At any time, all other electrode contacts record EEG signals, which are routed through the SM to the headbox and the amplifier. **b**. Removal of the stimulation ($T_0$) and switch-matrix artifacts (opening at $T_0$ – ms, closing at $T_0$ +300ms) for two example raw bipolar iEEG traces (red) with overlaid artifact-corrected (black) traces. **c**. Flowchart of iEEG signal preprocessing.

Stimulation electrical pulses had a bi-phasic square waveform with 0.5ms per phase and without inter-phase interval (1ms total). They were delivered over a range of intensities (0.2-12mA) in a bipolar stimulating electrode (two neighboring contacts on the same electrode lead). Each repeated stimulation block lasted one hour and included a five-minute non-stimulated baseline and two stimulation protocols. 1) The brain mapping protocol consisted of single pulse stimulation with a constant intensity of 3 mA, delivered to all bipolar electrode in grey matter. Each stimulation electrode was stimulated at least three times with an inter-stimulation interval of 4.5 s (+ jitter of a Gaussian distribution of standard deviation of 0.2 s). 2) The stimulation-response curve consisted of single pulse stimulations randomly repeated three times at one of 16 intensities ranging from 0.2 - 12 mA. This protocol was delivered in four stimulating electrodes in each participant, selected in priority in the healthy or epileptic hippocampus and/or amygdala, with two additional electrodes chosen randomly in neocortex. The total number of hourly blocks varied per patient over a range of 14-70 (Supplementary Table 2).

##### Signal preprocessing

Stimulation can induce sharp non-physiological artifacts in non-stimulating channels (Supplementary Fig. 1b). To smoothly remove these artifacts in the time-domain prior to any filtering, we located and removed the stimulation artifact by the TTL. Removal was done by kriging, which replaces the recorded signal by a linear trend between the average voltage before and after the artifact with super-imposed Gaussian noise that has equal variance as the variance calculated before and after the artifact. With our system, there are additional stimulation-related artifacts that were similarly removed: (i) Switchmatrix opening around 160-140 ms pre-stimulation, (ii) pre-stimulation peak around 25 ms pre-stimulation, and (iii) Switchmatrix closing around 300 ms post-stimulation (Supplementary Fig. 1a). These artifacts are located using a 'findpeak' function in the given time windows. For each stimulation trial and channel, the SM opening (5 ms), pre-stim peak (3 ms), stimulation (15 ms), and SM closing (5 ms) artifacts are removed by kriging. Each recorded signal was then bandpassed (0.5-200 Hz) and notch filtered (50 Hz and harmonics).

After filtering, signal rescaling was applied to normalize amplitudes across channels and mitigate impedance-related differences (Supplementary Fig. 1c). For each channel, a scaling factor was computed from the mean absolute amplitude during a 5-min baseline segment (wake, no stimulation, bandpass 0.5–30 Hz). Signals were first normalized by this channel-specific factor and then rescaled by a global constant to align the across-channel mean amplitude to a reference value (≈ 50 µV).

$$s_c = \text{mean}(|x_c(t)|)$$

$$s_{global} = \frac{50}{\frac{1}{C}\sum_{k=1}^{C} s_k}$$

$$x_c^{scaled}(t) = \frac{s_{global}}{s_c} \cdot x_c(t)$$

After rescaling, data was downsampled from a sampling frequency of 1024Hz to 500Hz.

##### Data curation

For each stimulation protocol, the data across blocks were segmented into epochs spanning from -1 second to 3 seconds and organized with the dimensions: channels x trials x 2000 (sampling frequency, sf = 500 Hz). The data were cleaned, addressing artifacts at three levels specific to channels and trials:

1. Any iEEG recording channel sharing at least one contact with the stimulation electrodes were excluded.
2. All iEEG recordings from previously stimulated electrodes were assessed for recovery by comparing the area-under-the-curve (AUC) of their baseline signal (from -500 ms to -50 ms before the current stimulus) against surrogate values and if their AUC exceeded the 99th percentile of surrogate values. This indicated recording channels that had not yet recovered from stimulations and may not record adequate signals.
3. All iEEG recordings containing outlier value during the baseline or post-stimulation response (Z-score > 8) were visually inspected and excluded if a technical artifact or epileptiform discharge was found.

##### Signal processing

Cortical responses elicited in individual stimulation trials exhibit variability in magnitude and shape (Supplementary Fig. 2). Prior studies of cortico-cortical evoked potentials (CCEPs) typically sought to mitigate this variability by studying the average responses across 5-50 individual trials time-locked to the stimulating pulse. These average CCEP classically recorded with pial electrodes reveal two negative peaks generally at 20-30 ms and 70- 300 ms post-stimulation when using strips or grids of electrodes apposed on the pia matter, but polarity and variable latency is problematic when using stereo-EEG[6,10–14]. Past works have relaxed these temporal constraints to accommodate delayed responses, and used several different characteristics of the average CCEP waveforms to quantify its magnitude, including its root mean square[6,10,12,15], area-under-the-curve[16–18], the early and/or late peak amplitude[3,13,14,16,17,19–21], or the peak-to-peak amplitude[18], each presenting general short-comings that are particularly relevant when seeking to quantify short- and long-latency single-trial responses.

$$AUC = \sum_{t=0s}^{1s} |x(t)|$$

$$P_{N1} = max_{10 \leq t \leq 50ms} \, |x(t)|$$

$$P_{N2} = max_{50 \leq t \leq 400ms} \, |x(t)|$$

$$P2P = x(t)_{max} - x(t)_{min} \text{ for } 0 \leq t \leq 1s$$

The area-under-the-curve is sensitive to offsets from zero (Supplementary Fig. 2). The quantification of the N1 and N2 peaks is contingent on their strict timing windows and the assumption of an immediate response with a distinct two-peak shape of a given polarity, which is not always present (Supplementary Fig. 2, all examples). The peak-to-peak amplitude is resilient to offsets but still relies on identifying peaks.

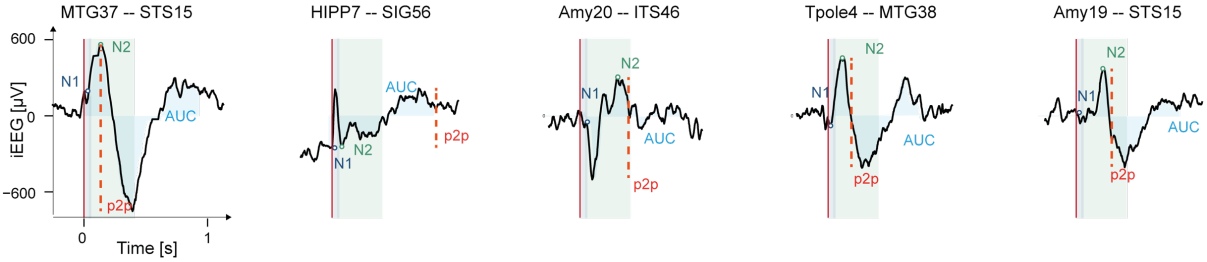


**Supplementary Figure 2 | Cortical response magnitude.** Examples of single-trial traces in five different connections, along with different metrics to measure the magnitude of the cortical response (cortico-cortical evoked potential, CCEP). Note the variability of evoked waveforms, their delay, their polarity and the problems incurred for different measurement methods. N1 and N2 peaks (polarity is ignored in stereo-EEG) and corresponding time windows are shown in blue and green, respectively. The peak-to-peak as orange dotted line, and area-under-the CCEP-curve as light blue area.

As a cornerstone of our work, we aimed to quantify cortical responses at the single-trial level, which required careful considerations regarding the reliability of methods to do so. We opted for the line-length transform (LL, Supplementary Fig. 3)[22] which is immune to offsets and polarity and avoids the need to identify pre-defined peaks. Indeed, the LL advantageously relies only one parameter, an integration window, over which the length of the absolute value of signal increments and decrements is summed:

$$LL(t) = \frac{\sum_{i=t-n}^{t} |x_i - x_{i-1}|}{n} * \frac{sf}{1000}$$

$$LL_p = max_{0 \leq t \leq 500ms}(LL(t))$$

where *LL(t)* is the line-length in ΔµV/ms calculated over a sliding window of 250ms (i.e. n=125 data-points at sf = 500Hz) at each post-stimulation timepoint between 0 and 500ms. LLp is the maximum line-length over that interval, and a Window of Interest (WOI) is centered at around LLp. Intuitively, line-length of the iEEG signal represents the amount of change in voltage per time, and likely reflects changes in neuronal firing rates underpinning cortical responses.

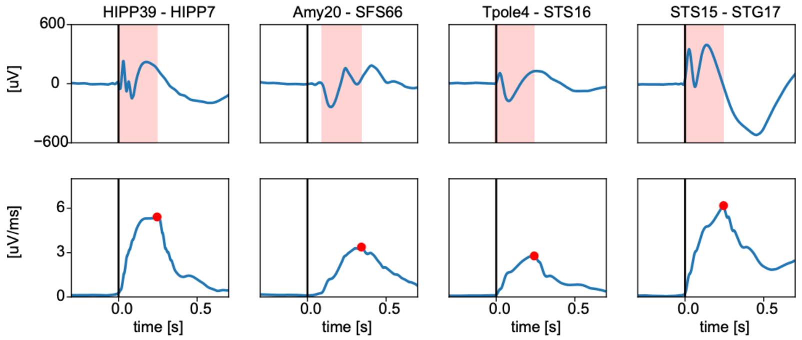


**Supplementary Figure 3. Line-length transform.** The mean cortical response from 195 trials in four connections (top) and the corresponding LL transform (bottom) over a 250ms sliding window (pink shading) are shown. The peak line-length is indicated as a red dot. Note that the cortical response in the SFS was delayed but still captured by our method.

We compared our line-length method to previously used metrics, finding that it correlated best with the peak-to-peak method, while circumventing the issue of defining peaks (Supplementary Fig. 4).

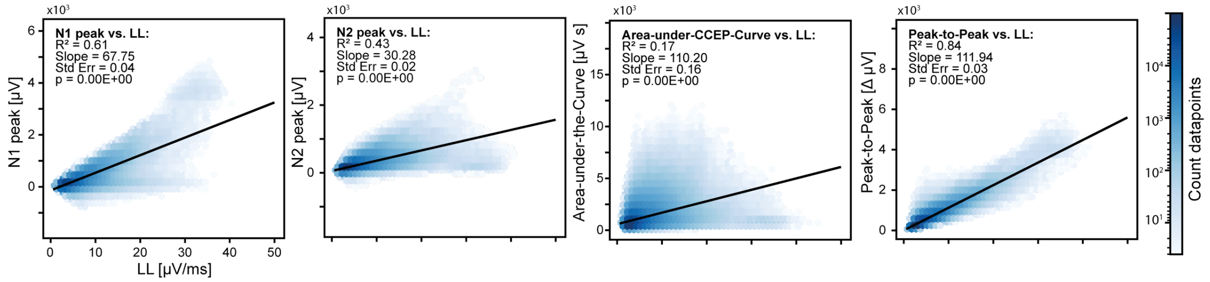


**Supplementary Figure 4 | Correlation analysis**. Correlation between linelength (LL) measurements from all significant trials (N = 3'409'378) and other metrics from left to right: N1 peak, N2 peak, area-under-the-curve and peak-to-peak. Hexagonal binning plot with a logarithmic color scale denotes the density of datapoints. A linear regression line (black) is overlaid and key statistical parameters—$R^2$, slope, standard error, and p-value are detailed in the upper left corner.

#### Effective connections

In this study we were interested time-varying signal flow within effective connections. Thus, we needed to develop a two-step method that could 1) establish the existence of an effective connection, accounting for the fact that signal flow may be inconsistent within that connection (Supplementary Fig. 5a-c1); 2) detect individual cortical responses of different magnitudes and waveforms (Supplementary Fig. 5a-c2). To account for this variability, we identified up to two potentially distinct cortical response waveforms for each connection. Technically, for each candidate stimulation-response pairs of electrodes, we derived two response signal cluster centers (CC) using an unsupervised clustering algorithm or rank two (K-means, with Pearson's correlation as the similarity metric). All the trials were clustered within a connection-specific window of interest (WOI) of 250 ms. The WOI is centered around the peak line-length ($LL_p$) of the trial-averaged signal, therefore covering the period of largest cortical response after stimulation. This step accounts for longer latency cortical responses up to a delay window between 250-500ms.

To test the presence of an effective connection statistically, we compared the line-length of the cluster centers to a null distribution drawn from 400 surrogate timeseries at non-stimulated time (baseline) and processed identically. An effective connection was deemed present, if LLp in at least one of the two cluster surpassed the $95^{th}$ ($p \leq 0.05$, FDR-corrected) of the null distribution. In non-connected stimulation-response electrode pairs, the two cluster centers were typically antiphase and of low magnitude not surpassing the null-distribution (column 3 in Supplementary Fig. 5b1). Thus, in our study, a connection was effective, if some signals were transmitted from the stimulating to the recording electrode, at least occasionally.

#### Single cortical responses

For each identified effective connection, we next assessed which of the single trials led to individual cortical responses. To that aim, we tested whether the individual post-stimulation signals had increased line length and resembled one of the extracted connection-specific CCs. Technically, each single-trial was cross-correlated[14] to each of the two CC in the WOI, and the maximal Pearson's coefficient (ρ ∈ [-1,1]) was retained. We then defined a compound metric (M) that multiplies the signed and squared ρ with the cortical response line-length, to weight it according to the resemblance to one CC.

$$\rho_{max} = \max[max_{\tau}\ (xcorr(CC1, signal)), max_{\tau}(xcorr(CC2, signal))]$$
$$M = sign(\rho_{max}) * \rho^2_{max} * LL$$

Where $\rho_{max}$ is the maximal cross-correlation (xcorr) between signal recorded at single-trial level and CC1 or CC2 within interval $\tau$, and $M$ the corresponding compound metric combining $LL$ and $\rho^2_{max}$ while conserving the sign of $\rho_{max}$. Intuitively, this approach quantifies the amount of line-length attributable to the expected response waveform. We then determined whether the value of $M$ was above that of a null distribution drawn from 400 non-stimulated baseline timeseries and computed identically ($p \leq 0.05$, without FDR-correction). Thus, in our study, cortical responses at single-trial level met two criteria: 1) the magnitude of the signal was increased compared to non-stimulated times, 2) the cortical response's waveform was similar to that of averaged trials.

#### Signaling latency

Different effective connections may signal with different latencies. Indeed, signaling latency could indicate the true 'synaptic distance' between two channels, that is how many synapses have to be passed until a cortical response is visible. For each connection, we calculated the 'peak latency' based on existing methods[2], that is the delay to the appearance of the first peak of the cortical response since stimulation. All trials with a significant cortical response were baseline corrected by subtracting the median of the signal voltage between -500:-20ms pre-stimulation. A statistical threshold was set to +/-3.4 standard deviations based on the baseline period of the baseline-corrected averaged signal. The latency of the first peak passing this threshold is considered the peak latency. Shorter and longer peak latencies likely represent direct (e.g. oligo-synaptic) and indirect (polysynaptic) connections[11] and this property is conserved across trials. In additional analyses, we calculated the peak latency for each brain state separately and found that they were conserved.

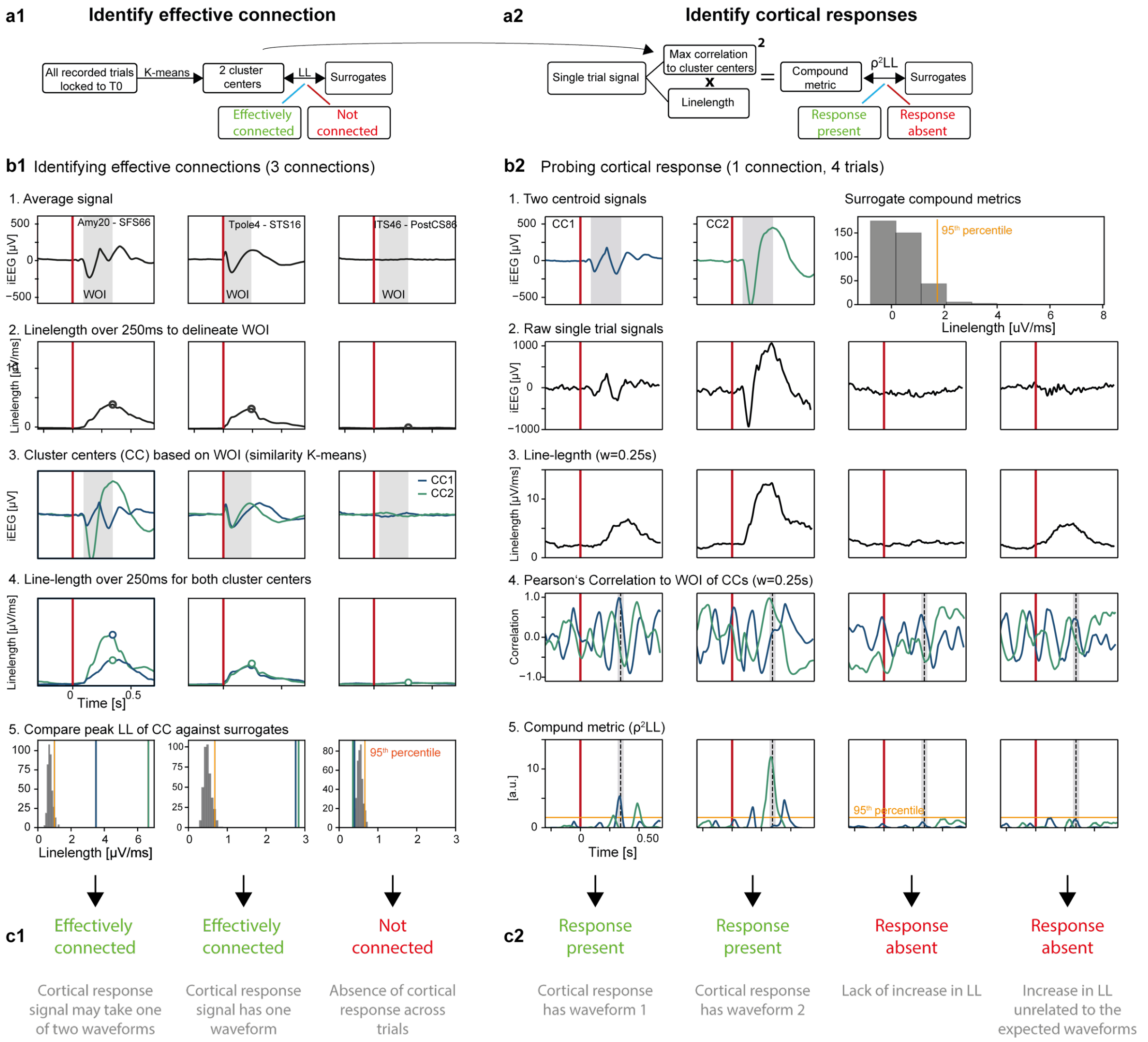


**Supplementary Figure 5 | Effective connections and cortical responses**. **a**. Flowchart of our methods for the detection of effective connections (a1), and of single-trial cortical responses (a2). **b1**. Detection of effective connections based on cluster average line-length (LL) to allow for variable waveform. For three illustrative connection examples (columns), the main steps to calculate the cluster centers from 195 single-trials are shown. 1) The average signal across trials. 2) The corresponding LL transform highlights the peak LL (black circle), which defines a Window of Interest (WOI, grey shadow in rows 1 and 3). 3) Two cluster centroids (CCs) derived by the K-means algorithm with Pearson's correlation within WOI as similarity metric. Note how clustering accommodates different (leftmost), similar (middle), or absent cortical responses (rightmost). 4) The corresponding LL transforms of the two cluster centroids with the peaks as circles. 5) Null distribution of peak LL values from surrogate data at non-stimulating times (grey, threshold at 95th percentile in orange) with true values as blue and green vertical lines (CCs from 4). **b2**: Detection of single-trial cortical responses based on a compound metric that captures its magnitude and expected waveform. 1) The two centroids from the effective connections are taken as template waveforms. 2) Four single-trial cortical responses in one connection (leftmost in b1) showing variable waveforms (from left to right): small-and large-magnitude response, lack of response, and unrelated oscillation. 3) Corresponding LL transform. 4) Cross-correlation (Pearson's coefficient ρ) between CC1 (blue) or CC2 (green, waveform template in WOI from 1) and the single-trial cortical response (from 2). 5) The compound metric M (sign-conserved $\rho^2$ * LL) with the corresponding threshold (orange line, 95th percentile of single-trial surrogates). Note how the compound metric increases for cortical responses that resemble one of the cluster centroid, but remains below threshold when a response is absent, if the line length increases for unrelated reasons (e.g. spindle rightmost) **c**: Outcome of statistical testing for the presence or absence of an effective connection (c1) or a single-trial cortical response (c2)

#### Signaling probability

Based on our statistical method to detect cortical responses at single trial level, we can evaluate the actual signal flow within an effective connection, captured as a signaling probability $P_{AB}$: the frequency at which a response in the recording electrode B is observed upon stimulation in the stimulating electrode A.

$$P_{AB} = \frac{n_{AB}}{N_A}$$

where $P_{AB}$ is the response probability in the effective connection directed from A→B, calculated as the ratio of observed significant single-trial cortical responses in electrode B $n_{AB}$ over the total number of fixed-intensity stimulations delivered in electrode A $N_A$. $P_{AB}$ ranges from 0 (stimulation in A never leads to a response in B) to 1 (every stimulation in A leads to a CCEP in B). Note that the case $P_{AB} = 0$ is equivalent to finding no effective connection between to candidate electrodes A→B, which does not exclude an effective connection B→A, in unidirectional connections. Thus, our measurement constitutes a probabilistic measure of the dynamics in a directed connection, based on trial-by-trial targeted probing and direct observation of cortical responses.

#### Signaling directionality

Given that often $P_{AB} \neq P_{BA}$ , we propose a novel directionality index (DI), devised to quantify the directional flow between two candidate electrode pairs.

$$DI_{AB} = \frac{P_{AB} - P_{BA}}{max(P_{AB}, P_{BA})}$$

where DI → 0 indicates bidirectional signaling ($P_{AB} \simeq P_{BA}$), DI → +1 a pure efference (solely sending) and DI → -1 a pure afference (solely receiving). Note that $DI_{BA}$ is the negative of $DI_{AB}$ and that |DI| = 0.5 already means doubling of signaling in one direction, with exponential increases for |DI| > 0.5. For instance, whether $P_{AB}$ is 40% or 80%, and $P_{BA}$ is 20% or 40%, the resulting DI would consistently be 0.5.

### Supplementary results

#### Data curation

During data curation across participants, we excluded a median [IQR] 0.13% [0.06, 0.21] due to pre-stimulation anomalies (e.g. epileptic spike), 3.7% [3.16, 5.40] due to stimulation artifact and 1.5% [0.92, 2.45] due to slow post-stimulation recovery.

#### Effective connectivity

Effective connectivity has been previously mapped as the *incidence* of effective connections between any two given Destrieux's cortical areas among the large F-tract dataset of 583 participants. This approach asks the question: *how many participant* have at least one effective connection among all participants whose iEEG implantation covers a given pair of cortical areas. We found a linear correlation (Pearson's $\rho = 0.47$, $p < 10^{-58}$) between the connection incidences found in our and the F-tract dataset (Supplementary Fig. 6a). In grouping connections according to the brain regions derived from the F-tract dataset, we found that effective connectivity was highest within brain regions also in our dataset (Supplementary Fig. 6b, diagonal).

We performed additional analysis to characterize the effective connectivity found in our dataset. We found a median [IQR] connection density of 58% [47%, 66%], using our method to detect effective connections. In addition, we confirmed like others[2,14] that the connection incidence decreased almost linearly (linear fit, $R^2 = 0.95$, $p < 10^{-39}$) as the inflated Euclidean distance increases between the stimulating and recording electrode (Supplementary Fig. 6c). The inflated Euclidean distance is the Euclidean distance between electrodes mapped to the inflated MNI brain.

#### Signaling latency

At inflated Euclidean distance of >100mm, the proportion of long-latency connections reaches a plateau and represent ~50% of all connections (Supplementary Fig. 6d). Peak signaling latency was also lowest within as compared to across brain regions (Supplementary Fig. 6e).

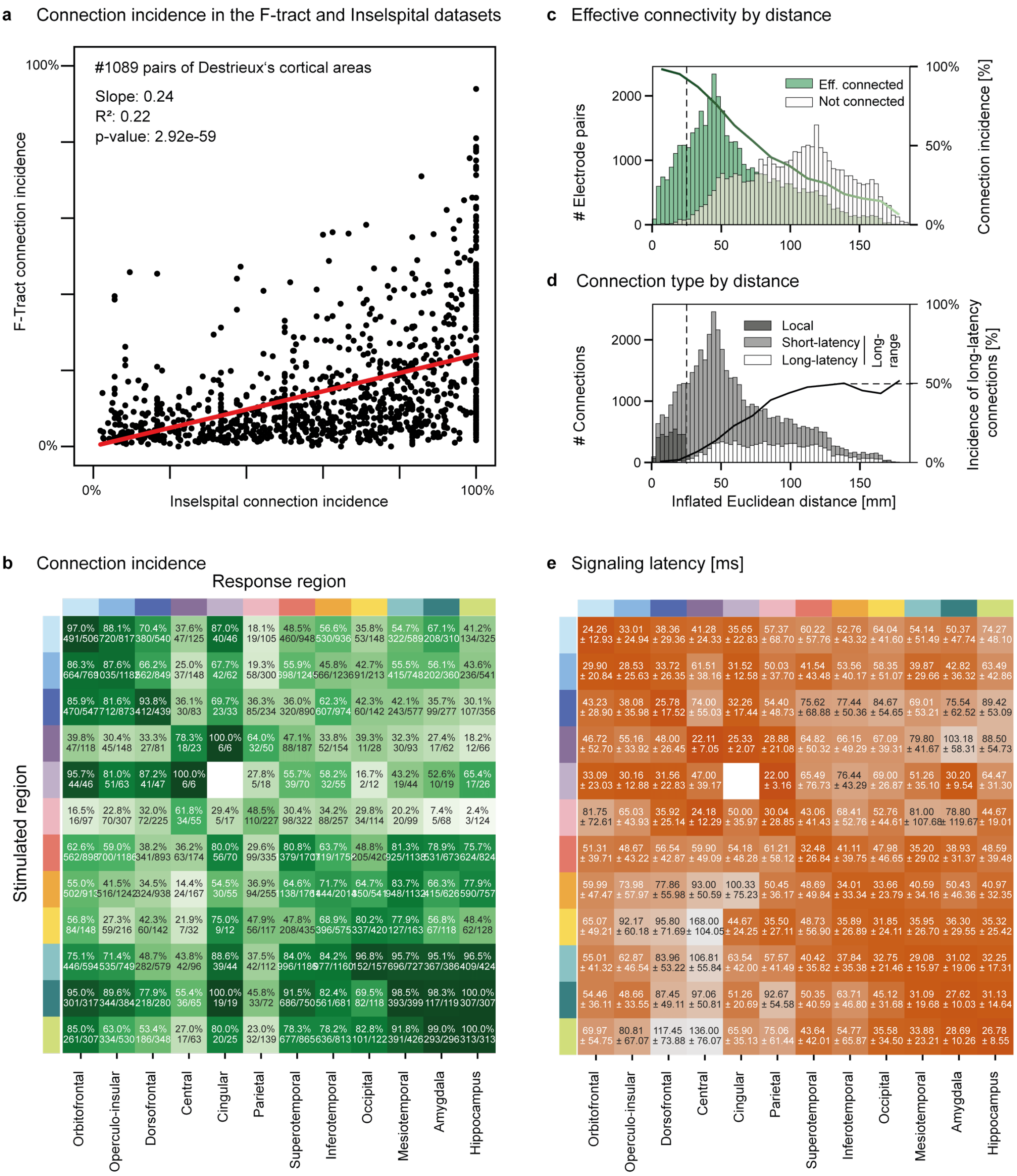


**Supplementary Figure 6 | Effective connectivity. a**. Mean connection incidence for pairs of cortical areas (Destrieux labels) in the F-Tract (N=583) and the Inselspital (N=15) datasets, showing a linear correlation (red line, Pearson's rho = 0.47, p< $10^{-58}$). **b**. Connection incidence across all possible intra-hemispheric pairs of brain regions in the Inselspital dataset (row: stimulation region, column: response region). **c**. Histogram showing the number of electrode pairs with (green) or without (white) an effective connection by inflated Euclidean distance in the Inselspital dataset. The dashed vertical black line delineates local from longer-range connections. The gradient-colored green curve represents the almost linear decay in connection incidence (%, y-axis on the right) as a function of inflated Euclidean distance between all tested electrode pairs (i.e. ratio green/white histograms). **d**. Stacked histogram showing the number of local (≤25 mm and same brain region, dark grey) versus long-range short-latency (≤65 ms, grey), and long-latency (>65 ms, white) connections by inflated Euclidean distance. The black curve represents the percentage long-latency connections out of the total effective connections. **e**. Corresponding mean [±SD] peak latency [ms].

#### Signaling probability

We performed additional analysis to characterize our method for measuring signaling probability. We visually evaluated in a number of connections, whether the method was indeed able to separate trials with and without cortical response. By averaging signals from trials classified as absent cortical response, we found mostly a flat line (Supplementary Fig. 7a). In a post-hoc analysis, we quantified this by evaluating the distribution of linelength from pre-stimulation signals as well as from signals with or without post-stimulation response (Supplementary Fig. 7b). We found that the distribution of linelength in non-responsive trials mostly resembled that of non-stimulated signals. Finally, we evaluated the consistency of signaling probabilities across participants. Supplementary Fig. 7c shows that the measured signaling probabilities for a given interregional connection were often very consistent across participants.

In a series of correlational analyses, we found that the signaling probability PAB and PBA were unrelated (Supplementary Fig. 8a). Further, signaling probabilities did not directly relate to the inflated Euclidean distance between stimulating and recording electrodes (Supplementary Fig. 8b), unlike connection incidence (Supplementary Fig. 6c). This indicated that although long-range connections may be rarer, they can signal with high probability. Also, we did not find that signaling probability directly related with the magnitude of cortical responses, suggesting that *how often* a cortical response is present, does not always relate to the average response magnitude (Supplementary Fig. 8b).

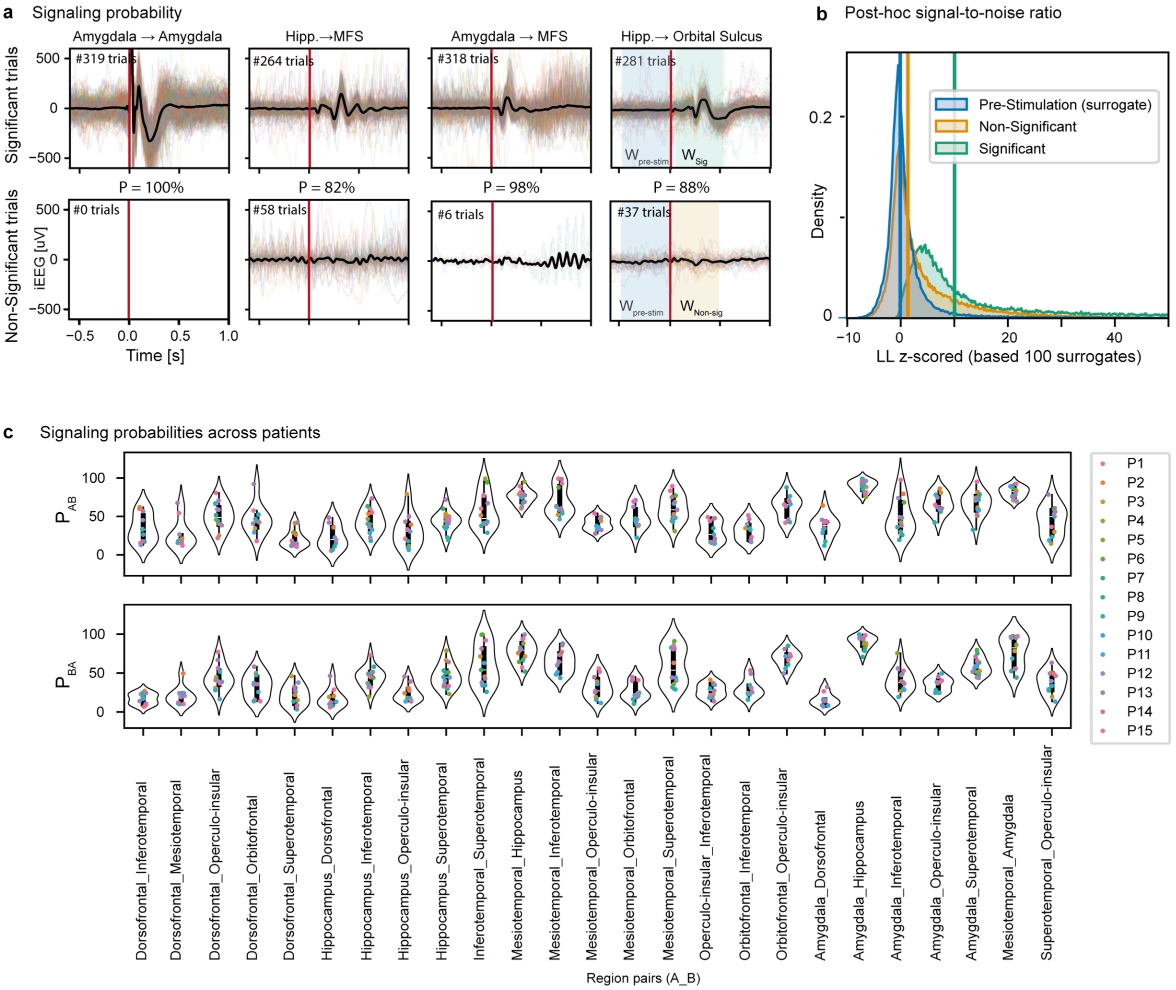


**Supplementary Figure 7 | Signaling probability. a**. Illustrative detection of single-trial cortical responses. The signaling probability (P) indicated above is calculated as the number of significant trials over the total number of stimulations. **b**. Post-hoc analysis of the distribution of the normalized linelength (z-scored based on surrogate distribution, interpretable as signal-to-noise ratio) for significant, non-significant and surrogate (i.e. non-stimulated) single-trial cortical responses. Colored vertical lines indicate the average of each distribution, which is largely superior for the trials deemed significant. **c**. Distribution of participant-averaged signaling probability in both directions (top: AB, bottom: BA) for pairs of brain regions covered in at least 7 participants.¬

#### Signaling directionality

We performed additional analysis to characterize our method for measuring signaling directionality. First, we compared our approach to directionality based on signaling probabilities to that of Entz et al., based on cortical response magnitudes in one and the other direction. We slightly modified the formula for the directionality index (DI) proposed by Entz et al., as follow:

$$oDI_Z = \frac{Z_{AB}}{Z_{BA}}$$

$$DI_Z = \frac{Z_{AB} - Z_{BA}}{max(Z_{AB}, Z_{BA})}$$

where $oDI_z$ is the original formulation of the DI by Entz et al. based on the ratio of Z-scores from 0 to +∞ and $DI_z$ is our modified formulation bound between -1 and +1 (see methods). This allowed us to compare our $DI_p$ based on probabilities with a $DI_z$ based on response Z-score magnitudes.

We found that the two measurements tended to agree when our $DI_P \rightarrow \pm1$, which makes intuitive sense, because if there is no effective connection in one or the other direction (i.e. unidirectional), this will be reflected equivalently in the very low response magnitude or probability (Supplementary Fig. 8a). However, for $DI_P \in [-0.9, 0.9]$, there was a very wide dispersion of $DI_Z$. Of specific concern, many connections in which we found a $DI_P \rightarrow 0$ (i.e. bidirectional), would have been attributed a $DI_Z \neq 0$ (dark vertical high density of points centered on $DI_P = 0$). Thus, $DI_P$ and $DI_Z$ are measuring different attributes of an effective connection, the former the frequency of signals transmitted in one and the other direction, the latter the relative signaling magnitude. For the study of signal flow, we must reject the measurement of directionality by comparing the magnitude of cortical responses.

Additionally, we found that our $DI_p$ was not influenced by the distance between the stimulating and recording electrodes (Supplementary Fig. 8b). We also asked whether a different signaling latency in one and the other direction could reflect directionality. However, we did not find a strong relationship between the delta peak latency and our $DI_p$ (Supplementary Fig. 8c).

All in all, correlational analyses in Supplementary Fig. 8 show that our dynamic measurements of signaling probabilities over time in one and the other direction and the resulting directionality index do not strongly correlate with average measurements such as the magnitude or latency of cortical responses. Also, long-range connections over a wide range of inflated Euclidean distance signaled with very variable probabilities and directionality, leaving the question of inter-regional specificity open.

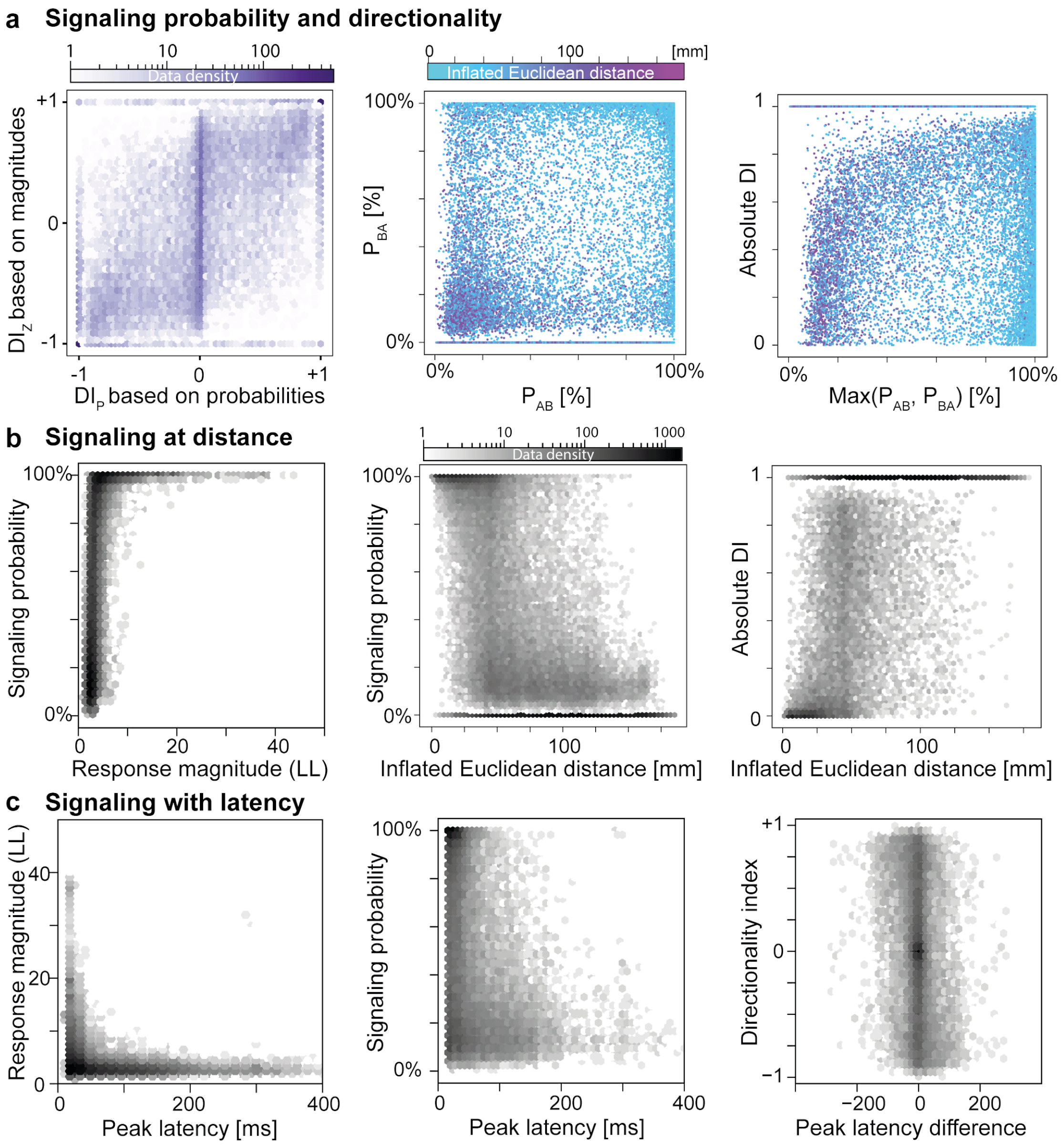


**Supplementary Figure 8 | Signaling probability, latency and directionality**. **a**. Definition of the directionality index based on the ratio of bi-directional probabilities: percentage significant responses in electrode B out of total number of stimulations delivered in electrode A, and inversely. Left: comparison of a directionality index based the ratio of cortical response probabilities ($DI_P$, our contribution) vs. magnitudes ($DI_Z$, possible alternative). While the two tend to agree when $DI_P \rightarrow \pm 1$, $DI_Z$ may take any value when $DI_P \in [-0.9, 0.9]$, ruling it out as a viable alternative. Middle: Each effective connection is represented by a datapoint placed according to its signaling probabilities in both directions ($P_{AB}$ vs. $P_{BA}$) and colored by the inflated Euclidean distance separating the recording (A or B) from the stimulation electrode (B or A). Middle, right: Note the lack of correlation between $P_{AB}$ and $P_{BA}$ or DI and max[$P_{AB}$, $P_{BA}$]. **b**. Lack of linear relationship between signaling probability or the corresponding DI versus response magnitude and inflated Euclidean distance. **c**. Lack of linear relationship between signaling magnitude, probability or directionality with signaling latency.

#### Connection excitability

We had previously shown that our Excitability index (ExI) can reflect the pharmacological up- or down-modulation of cortical excitability.[22,23] We here used the same protocol in a subset of stimulating electrodes to assess regional variability. Briefly, we measured the growth in the magnitude of cortical responses evoked over a range of increasing intensities from 0 to 12mA. The area under this stimulation-response curve represents the excitability index (Supplementary Fig. 9a-c).

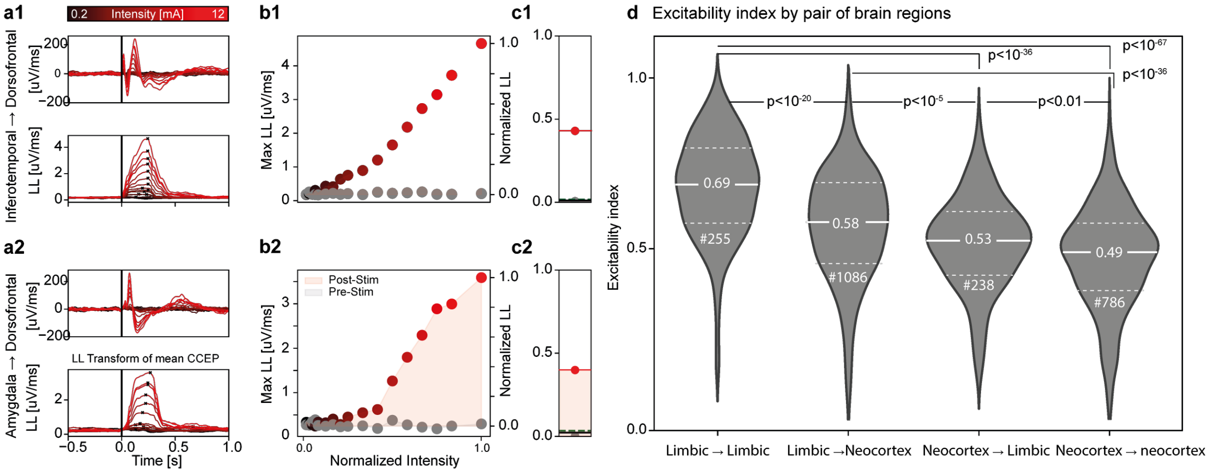


**Supplementary Figure 9 | Cortical excitability**. **a**. Two examples of stimulation-response curves and corresponding excitability index, each for one connection in one participant. The mean cortical response to the stimulation (top) and the corresponding linelength transform (LL, bottom) are shown with the stimulation intensity color-coded (0.2 to 12mA). **b**. Stimulation-response curves for stimulated (color) and non-stimulated surrogate signals (grey). **c**. The area-under-the-curve in b (pink) is the excitability index (ExI). **d**. Violin plot showing median and IQR distribution of the excitability index for inter-regional connections tested by Kruskal-Wallis ($p<10^{-77}$) and post-hoc Dunn test (# number of connections).

#### Cortico-limbic signaling

In our main analysis, we found that directionality was most marked in connections from the limbic region to all neocortical regions. This was corroborated by an analysis of excitability, that was strongest within limbic regions, and stronger in limbic efferences than afferences (Supplementary Fig. 9d).

Given that we pooled data from the two hemisphere and from epileptic and non-epileptic tissue in our main analysis, we here evaluated the data for a potential effect of laterality and epilepsy on our results. In our stratified analysis for laterality, we found that connections from the hippocampus or amygdala to neocortex were dominated by efferences in both hemispheres, but that this effect was significantly more marked in the right hemisphere as assessed by a two-way Kruskal-Wallis test for laterality (p=0.0034, left vs. right) with an interaction (p=0.014) for stimulated region (hippocampus vs. amygdala, Supplementary Fig. 10a). Given that less connections were sampled from the right hemisphere, the contribution of this stronger directionality in driving the main result is mild. More extensive analyses will be needed in the future, to confirm or infirm more pronounced cortico-limbic directionality in the right hemisphere.

In our stratified analysis for recordings from epileptic parenchyma, we did not find any significant difference in the probabilities or directionality (Supplementary Fig. 10b).

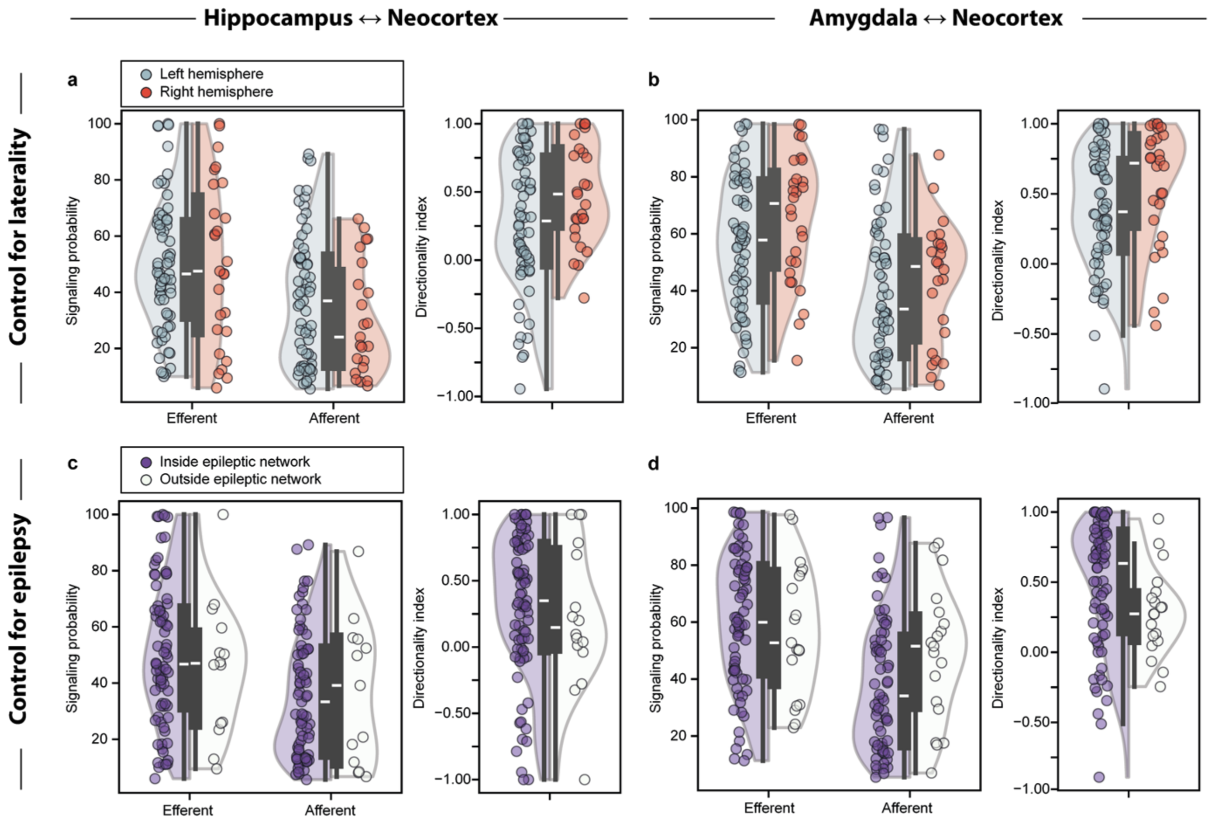


**Supplementary Figure 10 | Control analyses**. **a-b**. Control analysis for laterality, comparing connections between the hippocampus (a) or amygdala (b) with any ipsilateral neocortical brain region in the left or right hemisphere. Mean signaling probability and corresponding directionality index (DI) by region pairs across participant are shown as half-violin plots (left in grey, right in red). A significant increase in efferences was found in the right vs left hemisphere (p=0.0034) with an added effect of the stimulation region (amygdala vs. hippocampus, p= 0.014 for interaction, two-way Kruskal-Wallis test). **c-d**. Control analysis for epilepsy, comparing connections from and to epileptic (purple) versus non-epileptic (white) hippocampus (c, not involved in ictal discharges, occasional interictal discharges acceptable) or amygdala (d, not involved in ictal nor interictal discharges). No significant difference was observed in the probabilities and DI of epileptic vs. non-epileptic limbic structures (p> 0.05 for all, Kruskal-Wallis).

#### Cortico-cortical signaling

Although our focus was on cortico-limbic signaling, and our relatively low sampling of neocortical-neocortical connection does not allow firm conclusions, we here provide a comparative connectograms of all sampled connections (Supplementary Fig. 11). Three regions had a slight directional bias: orbitofrontal was receiving more than sending (PAB= 0.37 [0.18,0.63], DI= -0.12 [-0.48,0.06], p<0.01), whereas central (PAB=0.27 [0.21,0.62], DI=0.27 [0.03,0.58], p=0.02) and dorsofrontal (PAB=0.38 [0.18, 0.59], DI=0.08 [-0.20,0.57], p=0.01) were sending more than receiving, especially to orbitofrontal, operculo-insular and superotemporal regions. Although the remaining six neocortical regions did not show any average directional bias, they could have directionality in their connection to other specific regions. Thus, many effective connections imposed probabilities and directionality that were departing from the mean effect of distance in a consistent manner across participants. In addition to averages provided in the figures, a down-loadable software helps individual visualization of 42,435 connections recorded (Data availability).

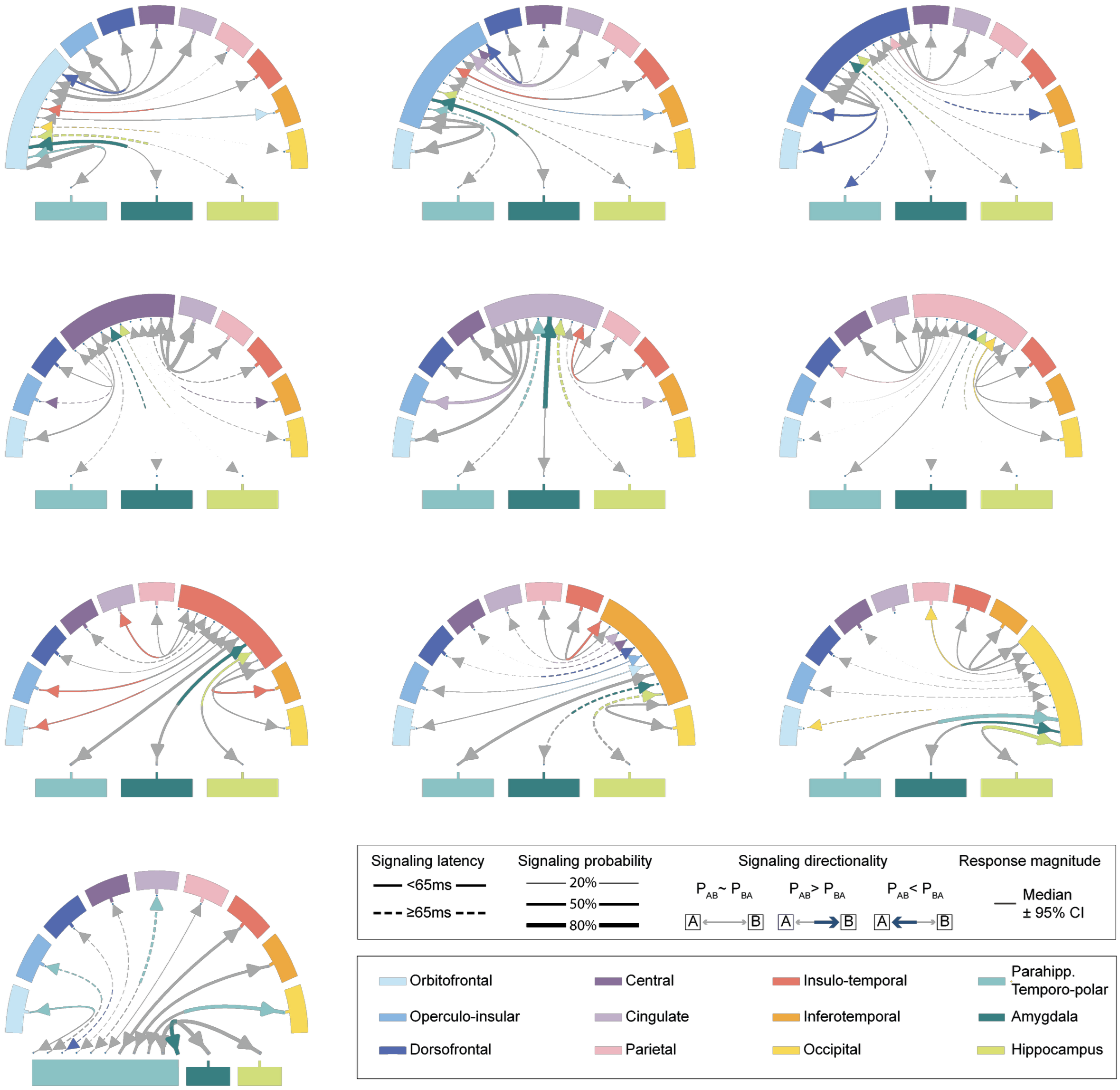


**Supplementary Figure 11 | Directional signaling across all brain regions.**
Connectograms showing all inter-regional average signaling probabilities (line width on the receiving end) and latency (dashed vs. full lines) between limbic structures and neocortical brain regions. For connections with a significant directionality index (post-hoc Wilcoxon test, $p < 0.05$), the direction with the higher signaling probability is highlighted with the color corresponding to the stimulated region. A graphical user interface can be downloaded at https://github.com/neuro-elab/EvM_Connectivity/releases/tag/Connecto_v0.1 to visualize signaling in the individual connections.

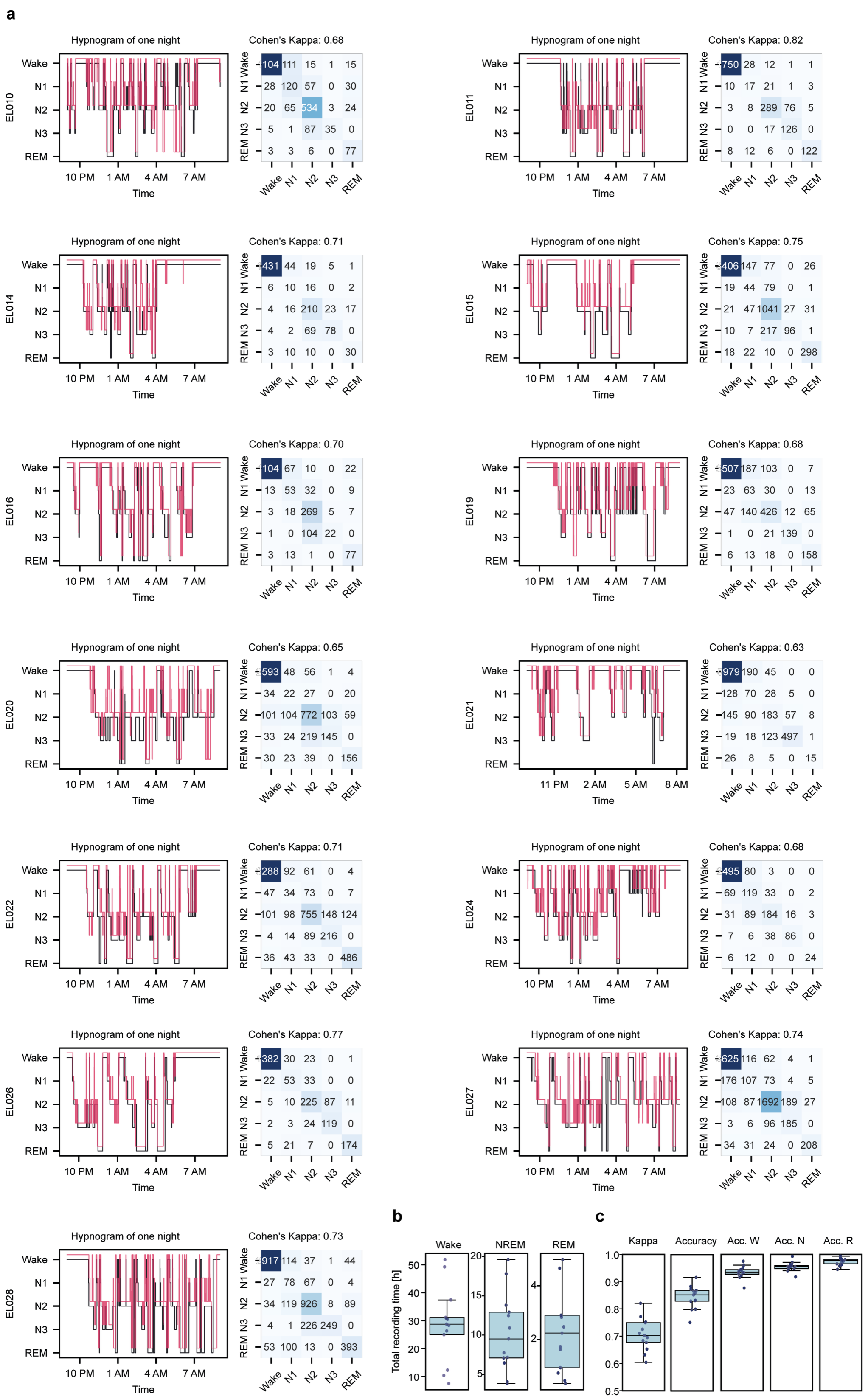


**Supplementary Figure 12 | Validation of visual sleep scoring**. **a**. For each participant, a hypnogram of 12h is shown that was scored by 2 independent scorers (black and red). The corresponding confusion matrix is plotted to the right. **b**. Total time (hours) in Wake, NREM (N2, N3), and REM across participants (data points). **c**. Sleep score agreement across participants (data points) based on Cohens' kappa, accuracy, and accuracy focused on wake, NREM (N2, N3), or REM.

#### Cortico-limbic signaling during sleep

Our final analysis was on potential modulation of signaling probability, directionality and connection excitability by NREM and REM sleep. We validated our scoring for vigilance stages in 13 participants with sufficient sleep data, by comparing the hypnograms obtained from two independent visual scorers (Supplementary Fig. 12).

We found in individual connection examples that the modulation by sleep of connections from the hippocampus or amygdala to the neocortex could be in opposite direction and was usually more marked for NREM sleep (Supplementary Fig. 13). Across connections though, the correlation between signaling probabilities during wake and NREM (Pearson's rho = 0.90) or REM sleep (rho = 0.95) were strong (Supplementary Fig. 13b). In a Bland-Altmann analysis for bias, we found the lack of overall bias (red dashed line in Supplementary Fig. 13b), but the connection that tended to have decreased probabilities outside of the confidence interval (blue dashed lines) in NREM and REM sleep were mostly hippocampal efferences. Upon detailed inspection of the probabilities by region-pairs, we found that efferences from the hippocampus to the cingulate gyrus, a major relay in Papez circuit was particularly decreased during NREM and REM sleep (Supplementary Fig. 13c). On the contrary, afferences to the hippocampus tended to slightly increase during NREM sleep (Supplementary Fig. 13c).

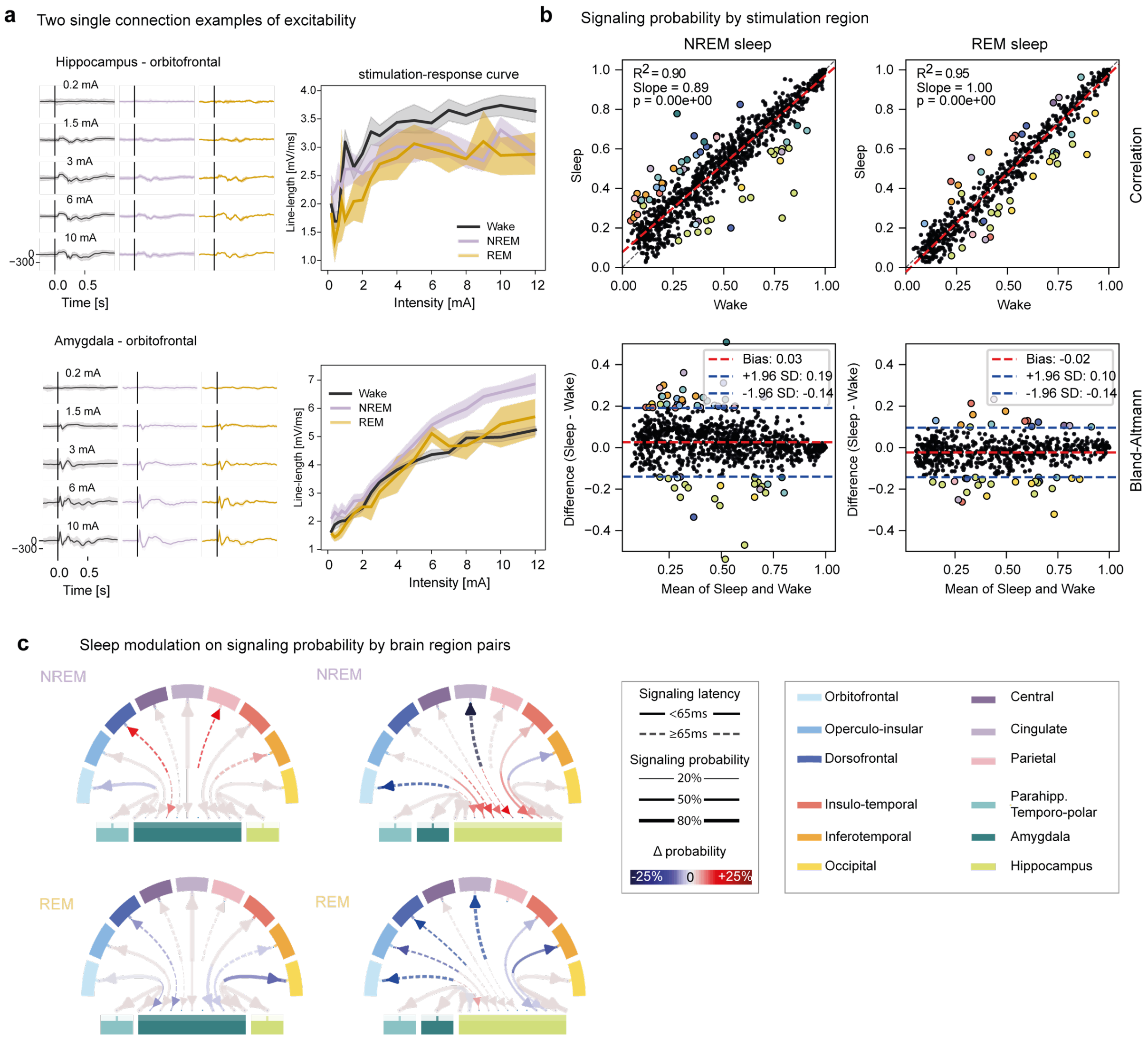


**Supplementary Figure 13 | Cortico-limbic signaling during sleep**. **a**. Two connection examples (stimulation in amygdala and hippocampus, response in orbitofrontal) illustrating gain curve during Wake, NREM and REM. **b**: top, mixed-effect linear model of the signaling probability during sleep compared to wake for each stimulated brain region (fixed effect) within participants (random effect). One dot is one region pair in one participant. Bottom: corresponding Bland Altmann plot showing the difference in signaling probability during sleep and wake against their mean value. Outliers are colored by the corresponding stimulation region. Note the high overall correlation and absent bias for sleep signaling probabilities compared to wake, with the exception of efferents from the hippocampus that tend to have decreased probabilities. **c**. Mean signaling probability between two brain regions for NREM and REM sleep. Colored arrows signify a significant difference in absolute signaling probability during sleep compared to wake (Δ probability, Wilcoxon test).
